# Effects of stimulus rate and periodicity on auditory cortical entrainment to continuous sounds

**DOI:** 10.1101/2022.09.04.506557

**Authors:** Sara Momtaz, Gavin M. Bidelman

## Abstract

The neural mechanisms underlying the exogenous coding and neural entrainment to rapid auditory stimuli have seen a recent surge of interest. However, few studies have characterized how parametric changes in stimulus presentation alter entrained responses. Applying inter-trial phase-locking (ITPL) and phase-locking value (PLV) analyses applied to high-density human electroencephalogram (EEG) data, we investigated the degree to which the brain entrains to speech vs. non-speech (i.e., click) sounds within and across tokens. Passive cortico-acoustic tracking was investigated in N=24 normal young adults utilizing EEG time-frequency and source analyses that isolated neural activity stemming from both auditory temporal cortices. We parametrically manipulated the rate and periodicity of repetitive, continuous speech and click stimuli to investigate how speed and jitter in ongoing sounds stream affect oscillatory entrainment. Both stimulus domains showed rightward hemisphere asymmetry in phase-locking strength with stronger and earlier responses to speech vs. clicks. Neuronal synchronization to speech was enhanced at 4.5 Hz (the putative universal rate of speech) and showed a differential pattern to that of clicks, particularly at higher rates. Phase-locking to speech decreased with increasing jitter but entrainment to speech remained superior to clicks. Surprisingly, click were invariant to periodicity manipulations. Our findings provide evidence that the brain’s neural entrainment to complex sounds is enhanced and more sensitized when processing speech relative to non-speech sounds. That this specialization is apparent even under passive listening suggests a priority of the auditory system for synchronizing to behaviorally-relevant signals.

## 1. Introduction

Temporal processing is a crucial component of auditory stream segregation and perceptual object representation (Picton, 2013). Temporal processing influences all levels of auditory skills ranging from sensory levels to higher cognitive processing including attention and memory (Allman et al., 2011; Toplak et al., 2006). Rhythm perception/production, entrainment, and time synchronization are subcategories of temporal processing in the auditory cognitive domain that vary between individuals (Grahn et al., 2012; Grondin, 2010; Thaut et al., 2005). In the broadest sense, entrainment refers to synchronization between two signals, which occurs by virtue of the phase-coupling (Cummins, 2009). Such rhythmic fluctuations in terms of brain-to-stimulus coupling are characterized by excitation–inhibition cycles of neuronal populations termed “neuronal oscillations” (Bishop, 1932; Buzsaki, 2006; Buzsaki et al., 2004; Lakatos et al., 2005). In the context of speech, oscillatory neural entrainment is the process by which auditory cortical activity precisely tracks and adjusts to modulations in the speech amplitude envelope via phase alignment between the neural and acoustic signal (Lakatos et al., 2019; Pikovsky et al., 2002). Entrainment is one of several important functions in auditory processing that can make communication seem effortless and automatic to healthy listeners and conversely, difficult in individuals with language learning disorders (Momtaz et al., 2021; Momtaz et al., 2022).

Entrainment applies to a wide range of physiologically important behaviors (Fujii et al., 2013; Herbst et al., 2016; Nozaradan et al., 2016) but is fundamental to speech (Peelle et al., 2012) and music (Fiveash et al., 2021) perception. For example, listeners must segment the continuous input speech signal into proper discrete units, which are then used as input for future decoding stages by auditory and non-auditory brain regions. In speech, the amplitude envelope’s rhythmic information reflects different aspects of sensory and motor processing such as segmentation, speech rate, and articulation place and manner (Peelle et al., 2012). Moreover, the perceptual system recovers the rhythmic structure of speech, which is important for spoken language comprehension (Poeppel et al., 2020). Sensory processing presumably benefits from neural entrainment as it could provide a temporal prediction mechanism to anticipate future auditory events before they arrive at the ear (Lakatos et al., 2013).

On the contrary, auditory signals like human speech might challenge an entrained system given its quasi-rhythmic temporal structure (Guenther, 2016; Levelt, 1993) and various rates speech can be conveyed to a listener. Still, speech production imparts temporal regularity to the signal envelope that is, on average, remarkably consistent in speed across languages (Poeppel et al., 2020). Indeed, the temporal syllabic rate of speech across various languages ranges from 2 to 8 Hz with a ‘characteristic’ periodicity of 4.5 Hz (Poeppel et al., 2020). It has been hypothesized that this range of rhythmicity is prioritized for speech perception and production (Assaneo et al., 2018). Moreover, speech intelligibility is optimal when the rhythmic structure of the signal falls inside a syllabic rate of 4-8 Hz (Doelling et al., 2014; Peelle et al., 2013). Thus, in addition to various speeds of representation, the quasi-periodic nature of speech effectively produces jitter which presumably also impact entrainment and subsequent auditory processing.

With regard to (a)periodicity, prior work has not fully elucidated how and to what extent aperiodicity might affect auditory processing (Breska et al., 2017; Doelling et al., 2015; Novembre et al., 2018). Some studies demonstrate comparable neuronal phase-locking patterns for both periodic and aperiodic non-speech stimuli (Breska et al., 2017; Morillon et al., 2016; Wilsch et al., 2015). However, upcoming events in speech and language can be predicted by non-periodic cues, e.g., based on syntactic or semantic features (Nguyen et al., 2015). Periodic stimuli, however, provide intrinsic predictability and subsequent computations that facilitate auditory perception and learning (Falk et al., 2017; Rimmele et al., 2018). Therefore, periodicity is probably fundamental in facilitating auditory processing because it provides temporal *predictability* to the system (Hovsepyan et al., 2020). At the very least, the brain must remain flexible and continuously adjust to changes in signal (a)periodicity to maintain robust processing. Yet, how different types of stimuli and the degree to which their aperiodicity affects auditory neural coding and entrainment remains unclear.

In the present study, we aimed to characterize how (i) rate, (ii) periodicity, and (iii) stimulus domain (i.e., speech vs. non-speech) affect auditory neural entrainment. In passive listening paradigms, we recorded multichannel EEGs in young, normal-hearing adults to assess neural entrainment to rapid auditory stimuli that parametrically varied in their speed (rate) and periodicity (temporal jitter). We analyzed the data at the source level to assess possible differences in hemispheric lateralization for entrained neural responses. The pacing of our rate manipulation assessed changes in neural oscillation strength for sounds presented slower than, at, and faster than the nominal syllabic rate of typical speech (i.e., 4.5 Hz). We reasoned that characterizing phase-locking strength across rates may demonstrate a preferred entrainment frequency of the system relative to rates that are considered to have special importance for speech perception. As a second manipulation, we evaluated the effects of signal periodicity on entrained brain activity. By adjusting the successive interstimulus interval between repeated tokens, we varied the stimulus delivery between fully aperiodic and periodic presentation. “Periodotopic” maps have been documented in temporal rate profiles from the inferior colliculus through the primary auditory cortex (Baumann et al., 2011; Langner, 2005). We thus extended this prior work by mapping periodicity sensitivity via oscillations of the scalp EEG. As a third aim, we assessed domain septicity of auditory neural entrainment. Studies using unintelligible sounds (Howard et al., 2010) have raised questions about whether entraining mechanisms reflect mere physical stimulus characteristics (Capilla et al., 2011) or higher-level functions unique to speech-language processing (Peelle et al., 2012). Thus, in addition to speech, we mapped rate and jitter functions non-speech (click) stimuli to test for possible domain specificity in entrainment strength. Neural responses were then compared to standard psychoacoustical assays of rate and periodicity sensitivity to assess the behavioral relevance of our EEG findings.

## 2. Methods

### 2.1. Participants

We recruited N=24 young adults (aged 20 to 39 years; 12 female, 12 male) to participate in the study. All participants had no history of neuropsychiatric illness and had normal hearing (i.e., air conduction thresholds ≤25 dB HL, screened from 500-4000 Hz; octave frequencies). History of music training, years of education, and handedness were documented. We required participants to have < 3 years of formal musical training since musicianship is known to enhance oscillatory EEG responses (Bidelman, 2017b; Trainor et al., 2009). All participants were monolingual English speakers and were right-handed (mean score at the Edinburgh handedness inventory = 79) (Oldfield, 1971). Participants gave written informed consent in compliance with a protocol approved by the Institutional Review Board at the University of Memphis and were monetarily compensated for their time.

### 2.2. Behavioral tasks and procedure

We used TMTFs and the CA-BAT paradigm to assess rate and periodicity sensitivity behaviorally, respectively (Bidelman et al., 2015; Harrison et al., 2018; Viemeister, 1979).

*Temporal modulation transfer functions (TMTFs)*. TMTFs are generally performed by modulating a carrier signal (e.g., noise) with a sinusoid at various rates and measuring the threshold modulation amplitude. TMTF is a psychoacoustic measure of listeners’ sensitivity to track amplitude modulations in an ongoing steady-state stimulus. The TMTF function describes amplitude detection thresholds (i.e., absolute sensitivity) as a function of modulation frequency (Bidelman et al., 2015; Dau et al., 1997; Viemeister, 1973; Viemeister, 1979). TMTFs were measured using a forced-choice, adaptive tracking task. Three consecutive 500 ms bursts of wide-band noise (100-10000 Hz) with 300 ms interstimulus interval (ISI) and 25 ms rise/fall ramping were presented binaurally using circumaural headphones (Sennheiser HD 280 Pro, 64 Ω). The noise was set at 74±1 dB SPL. The first and third noise bursts had no modulation; the second burst was modulated with a sinusoidal envelope at rates of 2.1, 3.3, 4.5, 8.5, and 14.9 Hz, identical to those used for the EEG recordings. Participants adjusted the degree of modulation imposed on the noise so that the fluctuation in the second noise burst was just detectable. Plotting the minimum detectable modulation depth across various carrier frequencies gives the TMTF, representing the I/O function across rates. Participants were allowed to adjust the modulation depth (measured in dB) using a slider bar on the computer screen until the difference between the target (modulated) and reference (unmodulated) intervals were no longer audible. The threshold was taken as the smallest modulation depth needed to just detect amplitude fluctuations in the stimulus. More negative thresholds reflect better task performance. This was repeated across rates to measure thresholds as a function of frequency. TMTFs were measured using the Auditory Interactivities Software (AI Core Team 2003).

*CA-BAT*. The computerized adaptive beat alignment test is a version of the beat alignment test which assessed participants’ behavioral sensitivity to periodicity, i.e., jitter (Harrison et al., 2018). The test consisted of 27 items lasting around 10 min that was presented about 74 dB SPL. Each item consisted of a beep track superimposed on a musical clip. The beep track alignment (d_r_) varied adaptively from trial to trial (0.5 ≤ d_r_ < 1) such that it was displaced in direction ahead or behind the music. Increasing d_r_ positions the beep-track closer to the musical beat and hence discrimination was harder. Critically, d_r_ was varied adaptively based on the listener’s trial-to-trial performance to converge onto their threshold for periodicity sensitivity. Participants were provided some sample music before the testing session as a training phase that includes instructions, audio demonstrations, and two practice questions. They were then given the 27 musical track test items in random order during the data collecting phase, with no item-to-item feedback. Each paradigm included a sequence of two-alternative forced-choice (2-AFC) trials. Two versions of a musical track were presented in each paradigm and their difference was in the overlaid metronome-like beep track. In one trial metronome and music were synchronized and in the other, the lure trial, they were displaced by a constant proportion of a beat. Participants were instructed to choose the one that was synchronized. The main output from the CA-BAT is an *ability score* (range −4 to 4), corresponding to the listener’s sensitivity to periodicity. A secondary output is an *ability_sem score*, corresponding to the standard error of measurement for the ability estimate. Both metrics are computed from the underlying item response theory model (Harrison et al., 2018). The paradigm was implemented in R (v.4.1.1) (R Team, 2013).

### 2.3. EEG recording procedures

*Stimuli*. EEGs were elicited using a click (100 µsec) and synthesized speech (60 ms) token /ba/. This speech token was selected as piloting testing determined it was the most identifiable token among several consonant-vowel options from previous neural oscillation studies (/ma/, /wa/, /va/, and /ba/ (Assaneo et al., 2018). In the rate experiment, each /ba/ stimulus was presented at 5 different rates (2.1, 3.3, 4.5, 8.5, 14.9 Hz). In the jitter experiment, this token was presented at the nominal syllabic rate of speech (4.5/sec) but we varied the trains’ periodicity by introducing random jitter in half the ISI between successive tokens. Jitter ranged from perfectly periodic (nominal ISI/2 ± 0% jitter) to aperiodic trains (nominal ISI/2 ± 80% jitter) in 5 equal steps from 0 to 80% (20% steps) (Krumbholz et al., 2003). Importantly, ISIs were uniformly sampled around the nominal rate (222 ms = 1/4.5) which maintained the overall average rate of stimuli between periodic and aperiodic conditions, allowing only the degree of periodicity to vary (**Figure 1**). Both click and speech stimuli were presented binaurally at 74.3 dB SPL via ER-2 insert earphones (Etymotic Research, Inc.). The 18 stimulus conditions (each N=1000 sweeps) were randomized for each participant.

**Figure 1.**
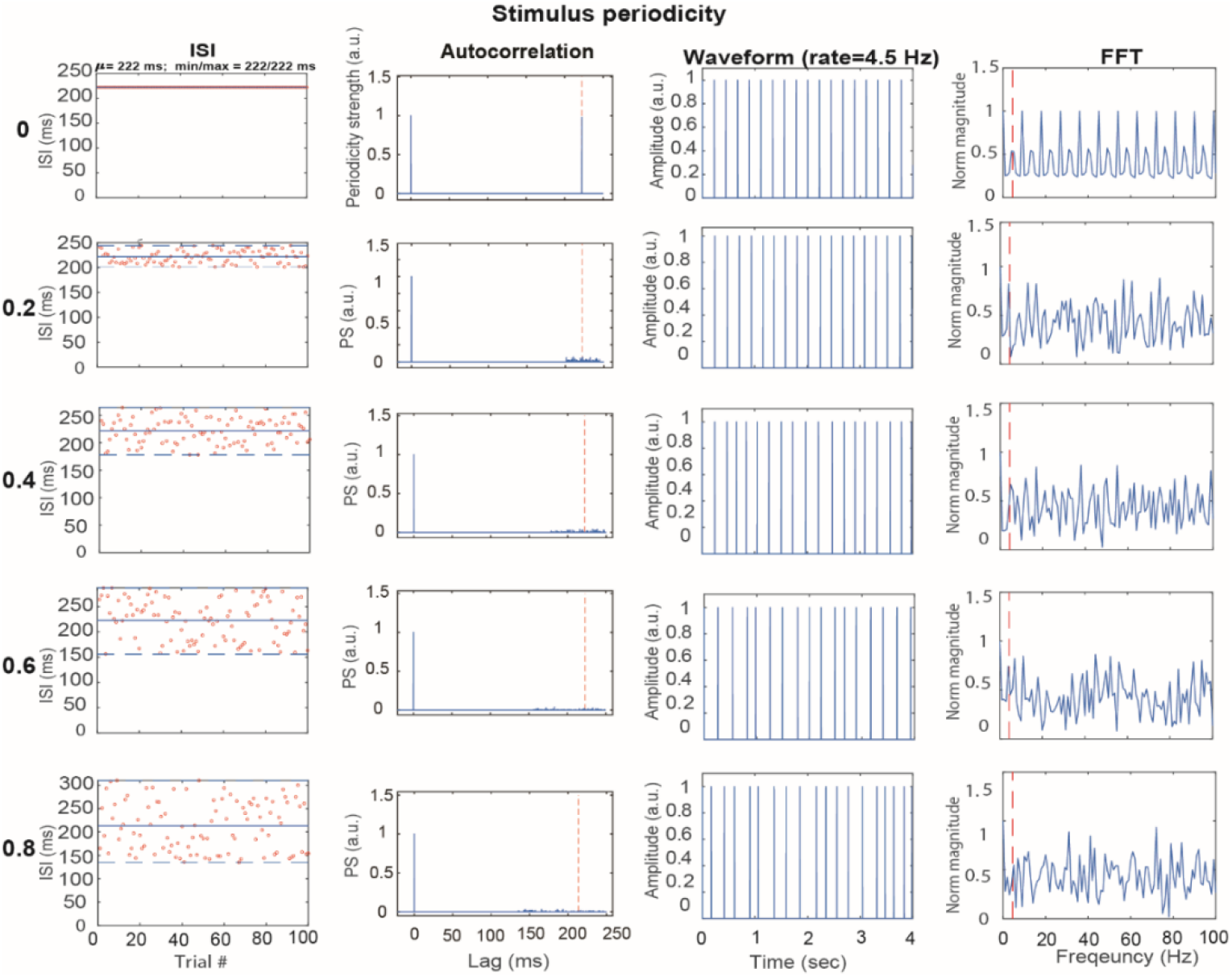
Acoustic properties illustrating the effects of (a)periodicity for the jitter stimuli (all 4.5 Hz rate with nominal ISI=1/4.5 Hz=222 ms. Half of ISI (111 ms) is spread in 20% stages from 0 to 80% ms (first column). The second column displays autocorrelation functions (ACFs), which show the degree of periodicity in the stimuli. Note the energy at 222 ms which becomes blurred around this nominal period with increasing jitter. (third column) Time waveforms. (fourth column) Fast Fourier transforms (FFT) as a function of jitter. As with the ACFs, not the decreased energy at 4.5 Hz with increasing stimulus jitter/aperiodicity. *EEG*. Participants were seated in an electrically shielded, sound-attenuating booth during EEG testing. They were asked to keep their eyes open and watch a silent selected movie (i.e., passive listening task). Continuous EEGs were recorded using a 64-channel electrode cap (Neuroscan QuikCap). Blink artifacts were monitored by placing electrodes on the outer canthi and superior and inferior orbit of the eyes. Electrode positions in the array followed the international 10-10 system (Oostenveld et al., 2001). Electrodes were maintained at <5 kΩ impedance during testing and were rehydrated halfway through the experiment as necessary. EEGs were recorded using Neuroscan Synamps RT amplifiers at a sample rate of 500 Hz. Data were re-referenced to the common average offline for subsequent analysis.

#### 2.3.1. Inter-trial phase-locking (ITPL) time-frequency analysis (within token analysis)

We used BESA Research 7.1 (BESA, GmbH) to preprocess the continuous EEG data for token-level time-frequency analysis (TFA). TFA was otherwise similar to our previous reports (Momtaz et al., 2022). Recordings were epoched into single trials from −10 ms to 56 ms and bandpass filtered from 10 to 2000 Hz (zero-phase Butterworth filters; slope = 48 dB/octave). Traces were then baseline corrected to ensure a zero mean pre-stimulus interval. Prior to TFA, the EEG data were cleaned from artifactual segments (e.g., blinks) by a 10 Hz high-pass filter. Paroxysmal electrodes were spherically spline interpolated. We then used a two-pronged approach for artifact rejection. Trials exceeding ±500 µV were first rejected using thresholding. This was followed by a gradient criterion, which was to discard epochs containing amplitude jumps of > 75 µV between any two consecutive samples.

Each listener’s single-trial scalp potential was transformed into source space using BESA’s Auditory Evoked Potential (AEP) virtual source montage (Bidelman, 2017a; Scherg et al., 2002). This applied a spatial filter to all electrodes that calculate their weighted contribution to the scalp recordings. We used a four-shell spherical volume conductor head model (Berg et al., 1994; Sarvas, 1987) with relative conductivities (1/Ωm) of 0.33, 0.33, 0.0042, and 1 for the head, scalp, skull, and cerebrospinal fluid, respectively, and compartment sizes of 85 mm (radius), 6 mm (thickness), 7 mm (thickness), and 1 mm (thickness) (Herdman et al., 2002; Picton et al., 1999). The AEP model includes 11 regional dipoles distributed across the brain including bilateral auditory cortex (AC) [Talairach coordinates (*x,y,z*; in mm): *left* = (−37, −18, 17) and *right* = (37, −18, 17)]. Regional sources consist of three dipoles describing current flow (units nAm) in the radial, tangential, and anterior-posterior planes. We extracted the time courses of the radial and tangential components for left and right AC sources as these orientations capture the majority of variance describing the auditory cortical ERPs (Picton et al., 1999). The two orientations were pooled for subsequent analyses. This approach allowed us to reduce each listener’s 64-channel data to 2 source channels describing neuronal currents localized to the left and right AC (Momtaz et al., 2021; Price et al., 2019).

We then performed a TFA on the source data to evaluate frequency-specific differences in neural oscillations between groups (Hoechstetter et al., 2004). From single-trial epochs, we computed a time-frequency transformation using a sliding window analysis (complex demodulation; Papp et al., 1977) and 10 ms/5 Hz resolution step sizes. These settings permit analysis of frequencies 10-80 Hz across the entire epoch window. The resulting spectral maps were produced by computing inter-trial phase-locking (ITPL) at each time-frequency point across single trials (Hoechstetter et al., 2004) according to Eq. 1:

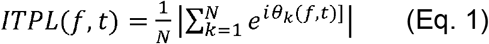

where *N* is the number of trials and *θ*_*k*_*(f,t)* is the phase in trial *k* at the time-frequency point *(f, t)*. The resulting spectral maps are 3D functions (time x frequency x ITPL), akin to neural spectrograms that visualize ITPL (phase-locking strength) rather than raw amplitudes. ITPL varies between 0-1 in which 0 reflects stochastic noise (i.e., absence of repeatable brain activity) and 1 reflects perfect trial-to-trial response repeatability (Tallon-Baudry et al., 1996). ITPL maps reflect the change in neural synchronization relative to baseline (−10 to 0 ms) and contain evoked neural activity that is time- and phase-locked to the eliciting repetitive stimulus (Bidelman, 2015; Cohen, 2014; Shahin et al., 2010). Maps were up-sampled by a factor of 10x (bicubic interpolated) in the time and frequency dimensions for further visualization and quantification.

To quantify the rate and jitter-related effects on auditory coding, we extracted the time course of the most robust frequency band from each spectrogram. We have typically observed dominant EEG energy to similar repetitive stimuli in the high β/low γ band frequency range (∼30 Hz) (Momtaz et al., 2021) (see also Fig. 3). Peak maximum amplitude and latency in the 30-40 Hz band from each band response were extracted automatically using MATLAB 2019 (The MathWorks, Inc).

#### 2.3.2. Phase-locking value (PLV) (across token analysis)

We computed phase-locking value (PLV) (Lachaux et al., 1999) between brain and stimulus waveforms to evaluate how neural oscillatory responses track speech and non-speech acoustic signals across different presentation speeds (rates) and periodicities (jitter). This complemented the within-token analysis (see Section 2.3.1) by allowing us to assess neuro-acoustic synchronization *across tokens* and at a longer temporal integration window of analysis (cf. sentence-vs. word-level processing). PLV was computed using the continuous EEG data. Mirroring our token-wise ITPL approach, we first transformed continuous EEGs into source waveforms (*SWF*) via matrix multiplication of the sensor data (*EEG*) with the AEP source montage’s dipole leadfield (*L*) matrix (i.e., SWF = L^-1^ x EEG) (Bidelman, 2018; Scherg et al., 2002). This resulted in two-time series representing current waveforms in source space projected from left and right AC. Source activity was then bandpass filtered (0.9-30 Hz) and 30 sec of continuous data extracted for submission to PLV analysis. Importantly, this yielded equal-length neural data per stimulus condition and listener. Identical processing was then applied to all rate and jitter conditions per participant.

We measured brain-to-stimulus synchronization as a function of frequency via phase-locking value (PLV) (Lachaux et al., 1999). Neural and acoustic stimulus signals were bandpass filtered (±0.5 Hz) around each nominal frequency bin from 1.1-30 Hz. PLV was then computed in each frequency band according to Eq. 2

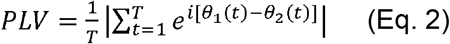

where □_*1*_*(t)* and □_*2*_*(t)* are the Hilbert phases of the EEG and corresponding evoking stimulus signal, respectively^1^. Intuitively, PLV describes the average phase difference (and by reciprocal, the correspondence) between the two signals. PLV ranges from 0-1, where 0 represents no (random) phase synchrony and 1 reflects perfect phase synchrony between signals. Repeating this procedure across frequencies (1.1-30 Hz; 0.3 Hz steps) resulted in a continuous function of PLV describing the degree of brain-to-stimulus synchronization across the bandwidth of interest (e.g., Assaneo et al., 2019b) (see **Figure 7**). We then measured PLV magnitude for each rate/jitter, stimulus type (speech vs. click), and participant. The magnitude was taken as the peak of each individual frequency-dependent PLV function within ±0.5 Hz of the nominal stimulus rate (see ▼s, **Figure 7**). Comparing PLV magnitude across increasing rates/jitters allowed us to characterize how brain-to-stimulus synchronization varied for speech vs. non-speech stimuli and between cerebral hemispheres. However, the omnibus ANOVA on PLV measures failed to reveal main or interaction effects with hemisphere (results reported below). Consequently, we collapsed LH and RH responses to focus on variations in peak PLV across stimulus manipulations (i.e., rates and (a)periodicities).

### 2.4. Statistical analysis

For EEG data, we used mixed-model ANOVAs in R (lme4 package) (Bates et al., 2014) to assess all dependent variables of interest. Fixed factors were hemisphere (2 levels, LH, RH), rate (2.1, 3.3, 4.5, 8.5, 14.9 Hz), periodicity (0, 20, 40, 60, 80% jitter), and stimulus domain (click, speech). Subjects served as a random effect. Based on the distribution of the data and initial diagnostics we transformed the data using a square root transformation. The significance level was set at α= 0.05. Tukey-Kramer adjustments were used for post hoc contrasts. Correlations (Pearson’s-r) were used to evaluate relationships between neural oscillations and behavior (e.g., rate ITPL vs. averaged TMTF; jitter ITPL vs. CA-BAT ability score).

## 3. Results

### 3.1. Behavioral data

*TMTFs (rate sensitivity)*. **Figure 2** shows the average TMTFs of our participants and the data of Viemeister (1973) for comparison. TMTFs show sensitivity (threshold) to amplitude modulation in wide-band noise measured at the five different rates. With increasing rate, participants showed better (i.e., more negative) detection thresholds corresponding to better sensitivity (i.e., temporal resolution). An ANOVA conduction on TMTF thresholds revealed a rate effect on TMTF thresholds [F_4,115_ = 10.44, p <0.001]. TMTFs thresholds typically worsen with increasing rates up to ∼100 Hz. However, at the low modulation rates used in this study—and consistent with prior psychoacoustic studies (Viemeister, 1973)—we find that rate sensitivity actually increases slightly between 2.1 and 14.9 Hz.

**Figure 2.**
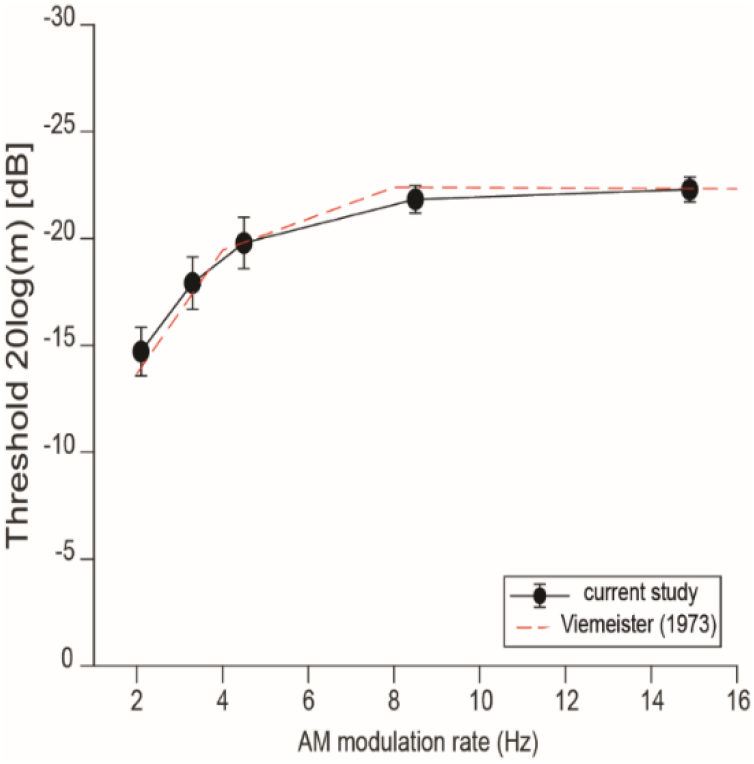
TMTF (temporal modulation transfer function) data showing behavioral rate sensitivity. TMTFs demonstrates temporal acuity for detecting amplitude fluctuations in continuous sounds as a function of modulation frequency. For very low frequencies <20 Hz, and consistent with Viemeister (1973), TMTFs demonstrate a high-pass filter shape, indicating slight improvements in behavioral rate sensitivity from 2 to 20 Hz. Error bars = ± 1 s.e.m.

*CA-BAT (aperiodicity sensitivity)*. The CA-BAT produced two different scores of periodicity sensitivity for each participant related to the absolute threshold (ability score) and its variance (ability_sem) (Harrison et al., 2018). Ability scores averaged 0.17 ± 0.19 (sem: 0.67 ± 0.04) across participants, consistent with prior psychoacoustic studies on jitter sensitivity (Harrison et al., 2018).

### 3.2. EEG oscillations within token (ITPL)

*ITPL magnitude*.**Figure 3** depicts ITPL spectrogram-like maps, reflecting evoked oscillatory responses localized to the auditory cortex (AC) that are phase-locked to the auditory stimuli. Visual inspection revealed 30-40 Hz spectrotemporal activity peaking ∼10-60 ms post-stimulation. Hence, we extracted the 30-40 Hz band time course to quantify differences in neural oscillatory representations across stimulus manipulations (rate, jitter), stimulus type (click, speech), and hemispheres (RH, LH) **Figure 4**.

**Figure 3.**
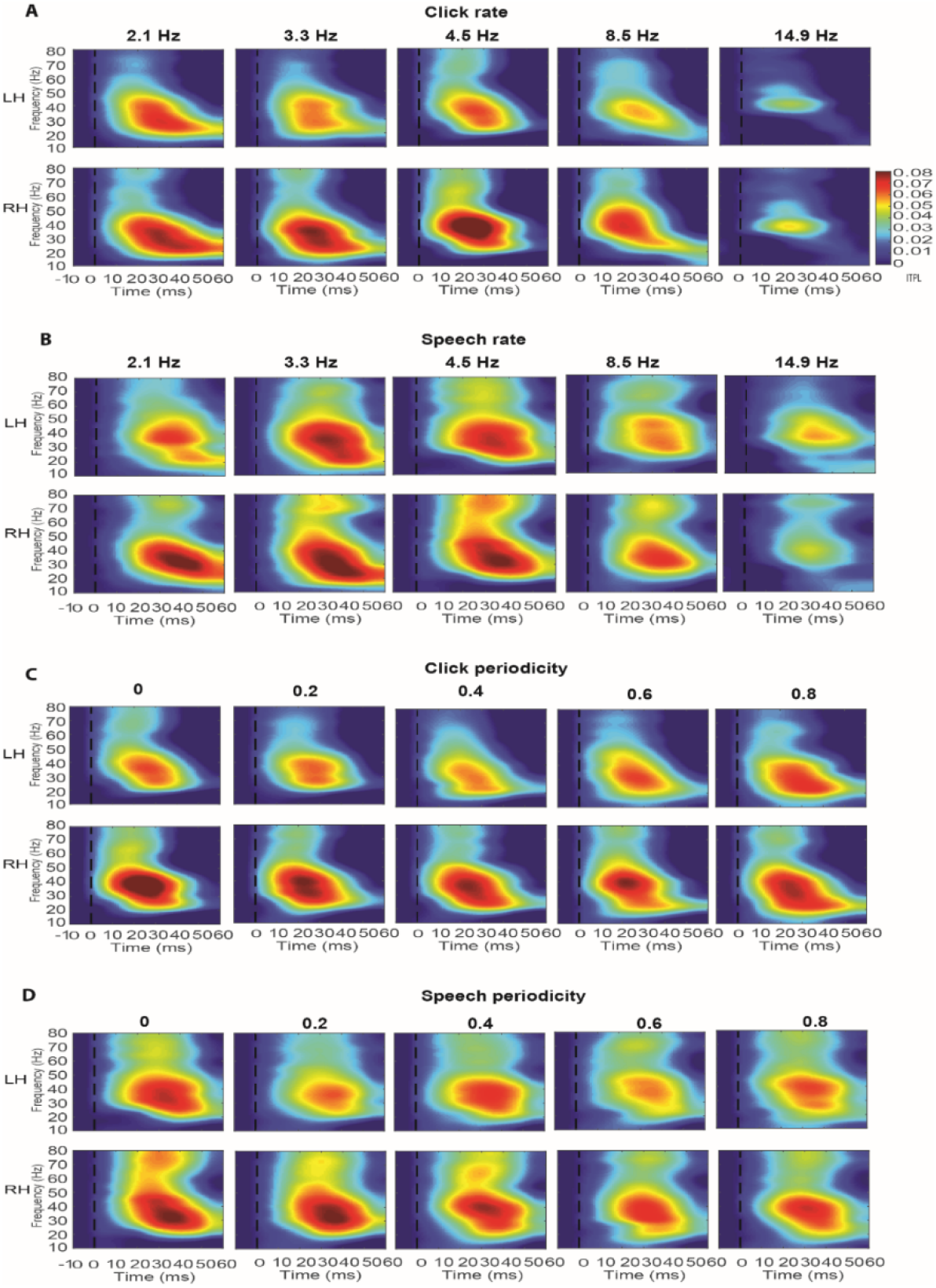
Grand average EEG ITPL spectrograms across rate and periodicity in speech and click. Time-frequency maps reflect neural oscillatory activity from the auditory cortex in each hemisphere. (A and C) click and (B and D) speech responses. ITPL maps depict “evoked” fluctuations of phase-locked EEG relative to baseline (i.e., the power spectrum of event-related brain potentials ERPs). t = 0, click stimulus onset. ITPL maps reveal robust neuronal synchrony, near the 30 - 40 Hz frequency region. Hotter colors indicate higher neuronal phase synchronization across trials. Left/right hemisphere, LH/RH.

**Figure 4.**
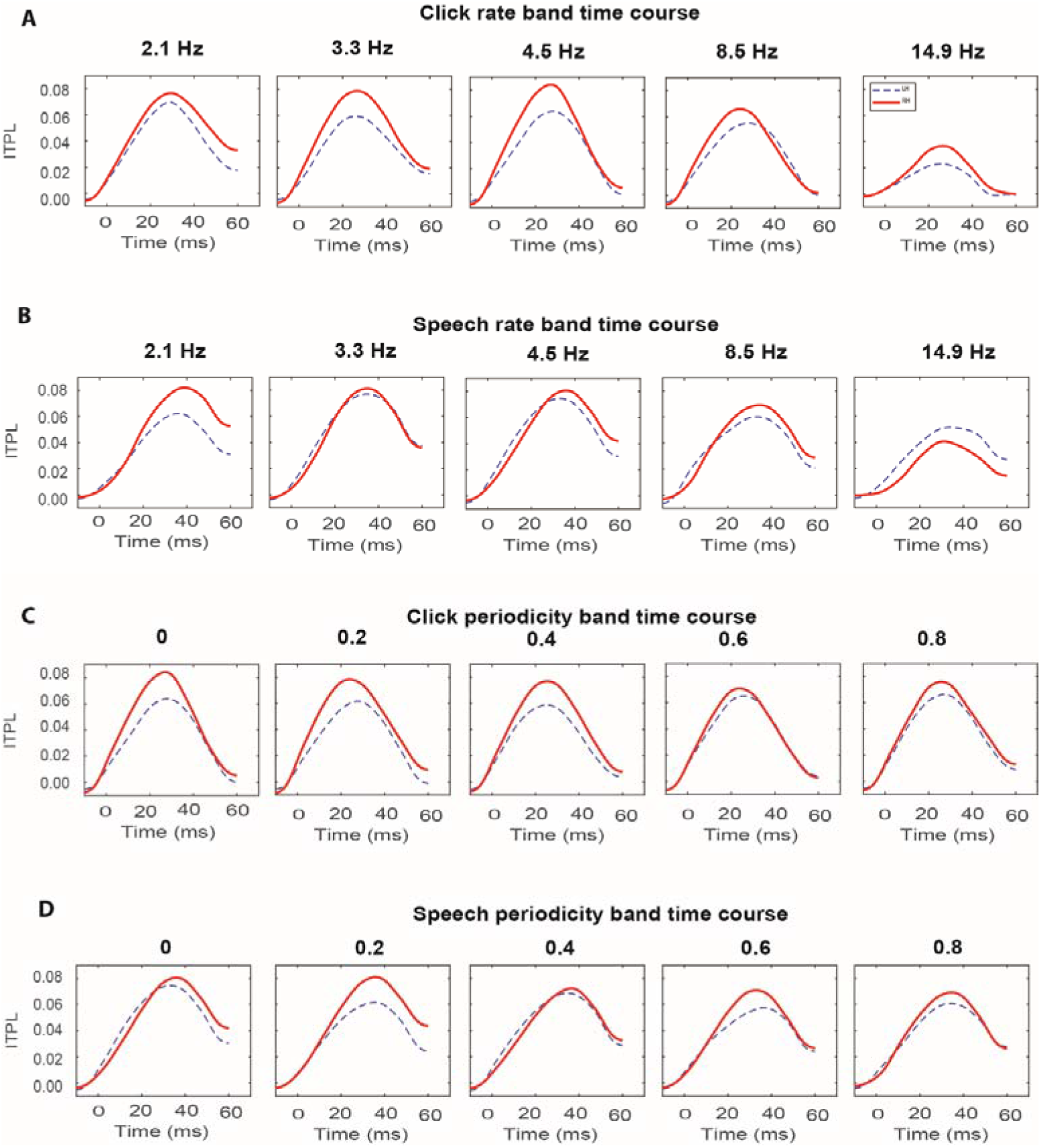
Grand average band time waveforms across rate and periodicity for speech and clicks. Time courses reflect temporal dynamics of phase-locking within the 30-40 Hz frequency band (see Fig. 3 ITPL maps). (A and C) click and (B and D) speech responses. Red solid lines= RH responses; blue dotted lines = LH responses.

For rate, an ANOVA on ITPL peak amplitudes revealed main effects of rate [*F*_*4,437*_ = 12.55, *p* < 0.0001] and hemisphere [*F*_*1,437*_ = 6.46, *p* = 0.01] on neural oscillation strength. No other main or interaction effects were significant. Planned Tukey-Kramer-adjusted contrasts revealed stronger amplitudes in 4.5 vs. 14.9 Hz for both click (*p* < 0.0001) and speech (*p* = 0.0009) stimuli (**Figure 5**). Post-hoc multiple comparisons of the hemisphere effect revealed lower LH phase-locking compared to RH regardless of the stimulus domain (i.e., speech vs. clicks).

**Figure 5.**
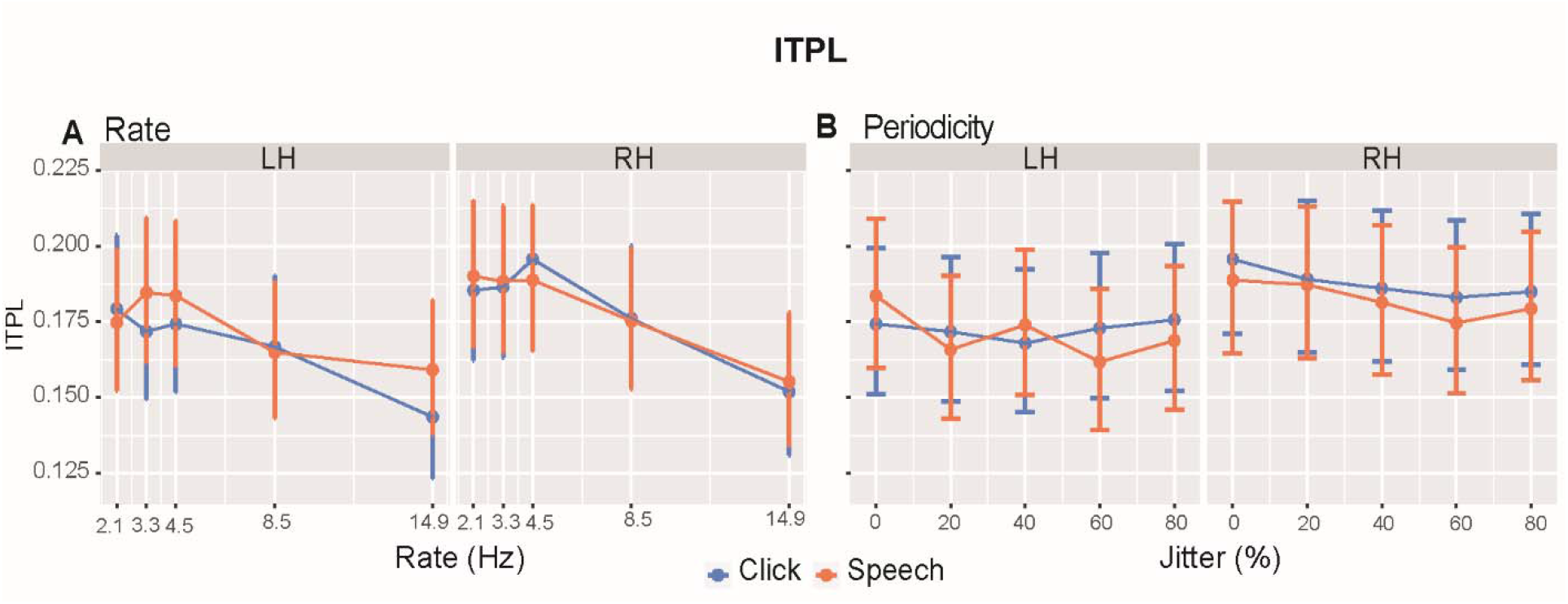
ITPL (within token) strength across rate and jitter. (A) Rate results. ITPL strength declines with increasing rate. Both clicks and speech stimuli showed increased ITPL strength in the RH vs. LH. (B) Periodicity results. ITPL was stronger for RH vs. LH for both speech and click stimuli. No interactions were significant. error bars = ±0.95 CI.

For periodicity, an ANOVA only showed a hemisphere effect [*F*_*1,437*_ = 14.98, *p* = 0.0001]. Again, RH phase-locking was more robust than LH for both stimulus types. No other main or interaction effects with ITPL strength were observed for periodicity.

#### ITPL Latency

Latency of the 30-40 Hz band response for the *rate* manipulation revealed main effects of rate [*F*_*4,437*_ = 4.06, *p* = 0.003], stimulus [*F*_*1,437*_ = 62.20, *p* < 0.0001], and a rate x hemisphere interaction [*F*_*4,437*_ = 2.73, *p* = 0.02] (**Figure 6**). Overall, the interaction was attributable to LH having delay responses compared to RH at 3.3 Hz in speech (p = 0.005). Separate analyses of RH and LH revealed response latencies were largely invariant to rate within each individual hemisphere (all *p*-values > 0.1).

**Figure 6.**
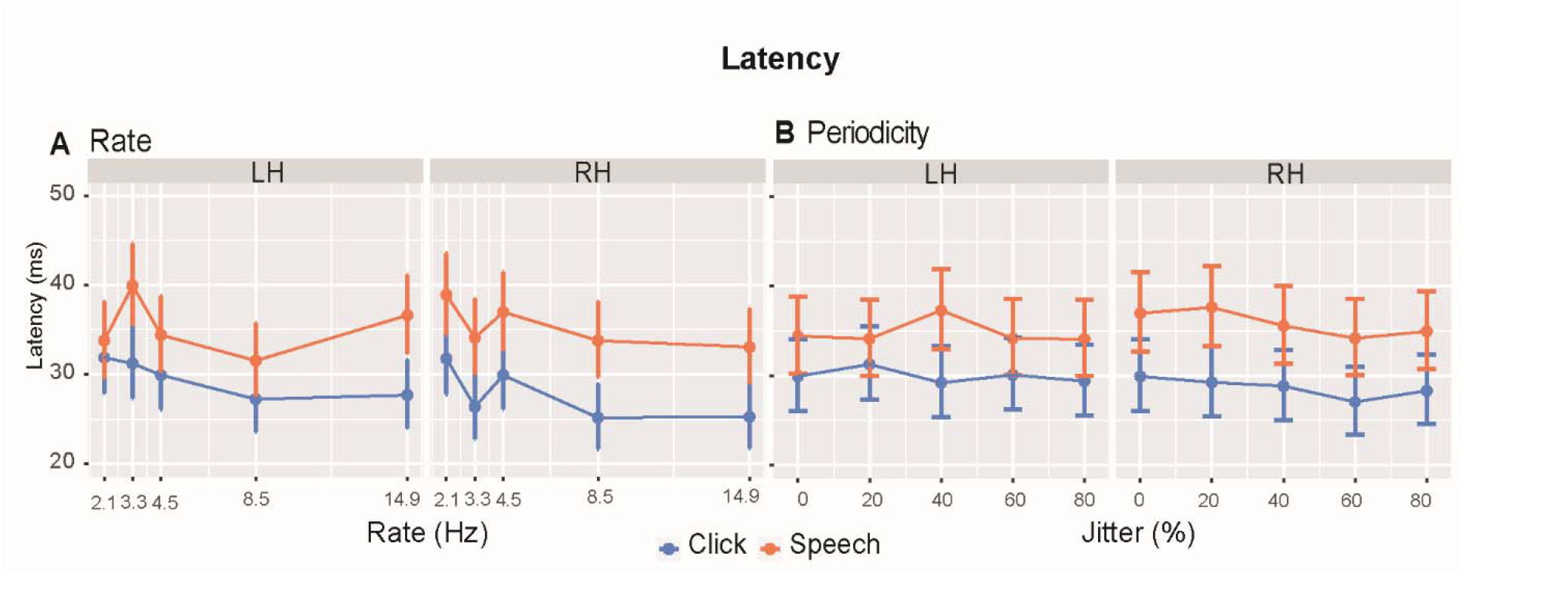
ITPL latency across rate and jitter. (A) Rate results. Latencies changed with rate, stimuli, and a rate x hemisphere interaction. Speech stimuli showed increased ITPL latency in both hemispheres. (B) Periodicity results. Responses were later for speech vs. click in both hemispheres. error bars = ±0.95 CI.

Analysis of *periodicity* on latency measures only revealed a stimulus effect [*F*_*1,437*_ = 49.39, *p* < 0.0001]. This was attributable to clicks producing faster responses than speech sounds across the board.

### 3.3. EEG oscillations across token (PLV)

PLV examined neural phase-locking *across tokens* and how the brain entrains over the entire stream of stimulus time. Raw PLV response functions illustrating changes in phase-locking strength as a function of frequency, stimulus manipulations (rate, jitter), hemispheres (RH, LH), and stimulus type (click, speech) are shown in **Figure 7**. Peak quantification of the PLV functions is shown in **Figure. 8**. In general, neural responses closely followed the speed of the auditory stimuli, showing stark increases in PLV at the fundamental rate of presentation as well as harmonically-related frequencies. PLV strength also varied for the 4.5 Hz stream with changes in jitter; more aperiodic sounds produced weaker neural entrainment.

**Figure 7.**
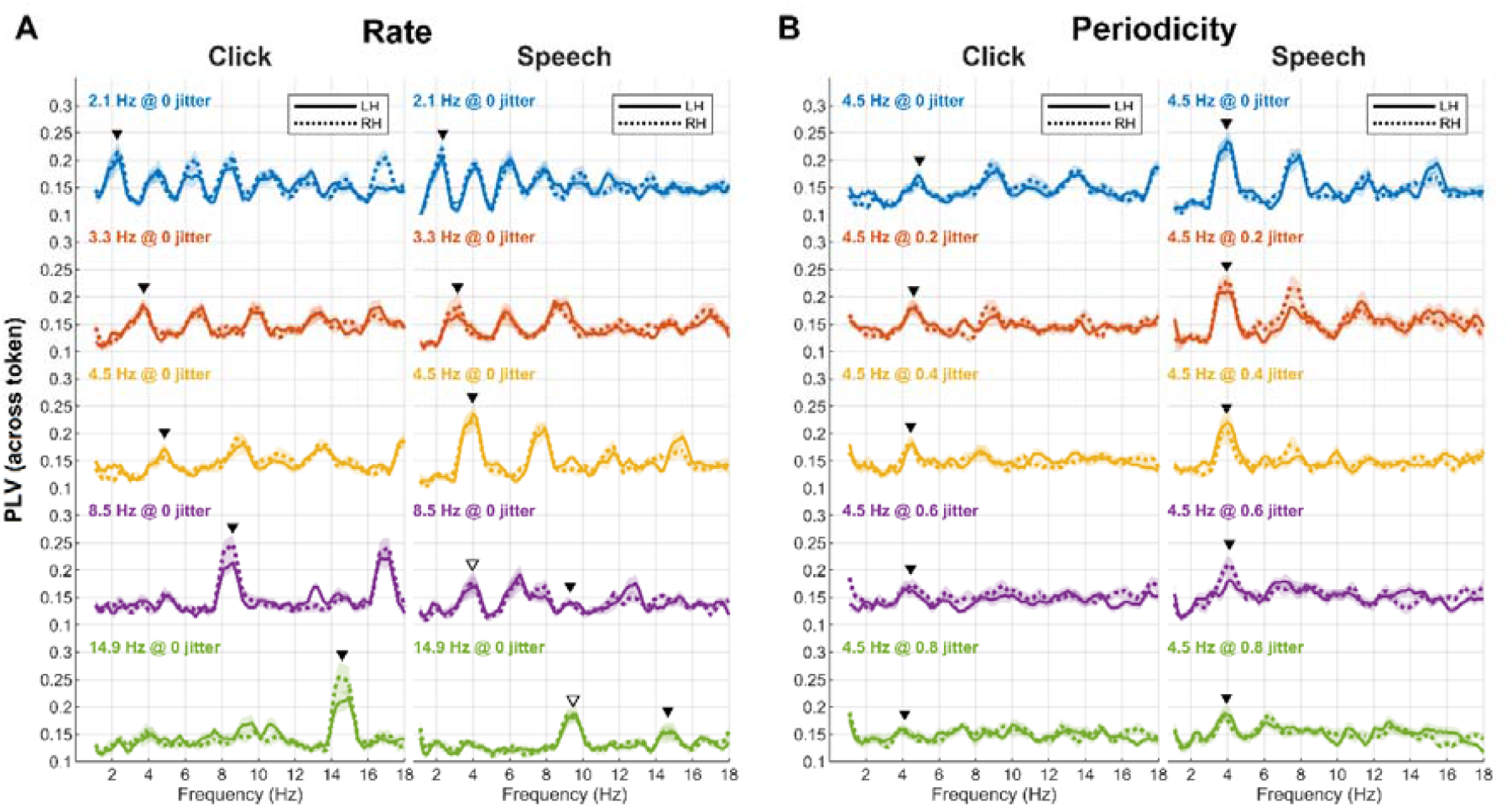
PLV (across tokens) as a function of rate, jitter, stimulus, and hemisphere. (A) Rate results. (B) Periodicity results. PLV showed enhanced activity at each fundamental frequency (▾) and integer-related harmonics. White triangles= subharmonics. Right and left hemispheres exhibited comparable responses.

**Figure 8.**
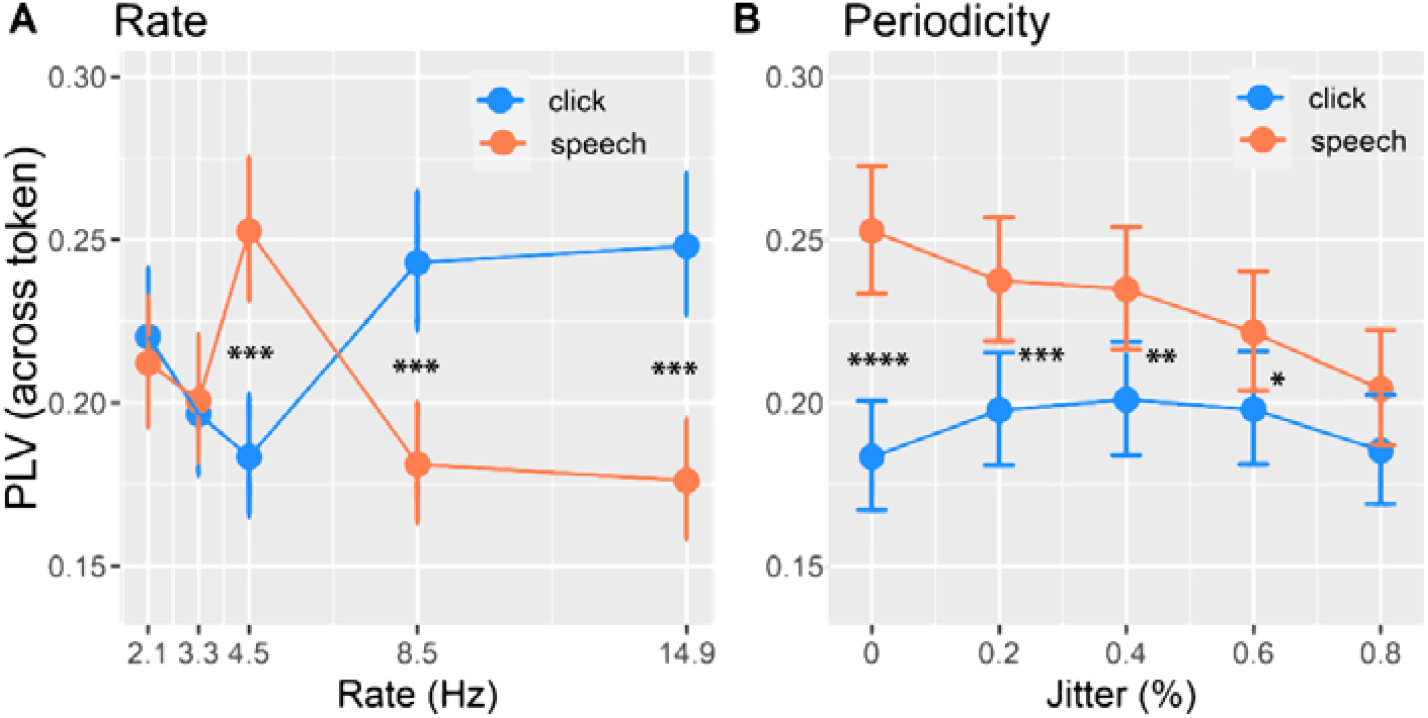
Peak PLV across rate and jitter. (A) Rate effects show a rate x stimulus interaction. Speech elicited increased neural entrainment at 4.5 Hz. (B) Periodicity effects show a stimulus x jitter interaction. Neural entrainment to speech is initially enhanced compared to clicks at low jitters but declines within increasing aperiodicity. PLV strength is invariant to increasing jitter for clicks. PLV = Phase locking value. error bars = ±0.95 CI. \**p*<0.05, \*\**p*<0.01 \*\*\**p*<0.001.

An ANOVA assessing rate effects on peak PLV showed a rate x stimulus interaction [*F*_*4,437*_ = 16.68, *p* < 0.0001] (**Fig. 8A**). Multiple comparisons showed stronger PLV for speech vs. clicks at 4.5 Hz (*p* < 0.0001). This pattern reversed at higher rates with speech eliciting lower PLV than clicks at 8.5 (*p* < 0.0001) and 14.9 Hz (*p* < 0.0001). These results suggest neural entrainment for speech is enhanced relative to non-speech signals specifically at 4.5 Hz and weakens precipitously for higher rates beyond what typically occurs in normal speech production.

For periodicity, we found a periodicity x stimulus interaction [*F*_*4,437*_ = 2.63, *p* = 0.03] (**Fig. 8B**). Except for 80% jitter, click vs. speech contrasts revealed higher PLV for speech across most jitter levels including 0% (*p* < 0.0001), 20% (*p* = 0.001), 40% (*p* = 0.005), 60% (*p* = 0.04). In other words, the overall pattern for speech exhibited a linear decline in which PLV strength decreased with increasing jitter. In contrast, entrainment to clicks remained relatively constant with increasing jitter. This interaction suggests that speech produces stronger neural entrainment across tokens than non-speech sounds and is also more impervious to disruptions in periodicity (i.e., signal jitter).

### 3.4. Brain-behavior relationships

We used correlations to explore the correspondence between neural responses and behavioral measures. TMTFs showed no correlation with within-token ITPL amplitude or latency metrics (**Figure 9**). However, when considering across-token entrainment, the average PLV across rates was highly correlated with TMTF thresholds (r = 0.56, p = 0.004); larger neural PLV was associated with poorer (less negative) behavioral thresholds. Finally, for the CA-BAT test, we did not find a relationship between the degree of periodicity sensitivity and any of the neural response measures (all p-values > 0.81). These findings indicate participants’ behavioral sensitivity to periodicity is not associated with their brain responses, which may be related to the active nature of our behavioral tasks vs. passive nature of our brain recordings.

**Figure 9.**
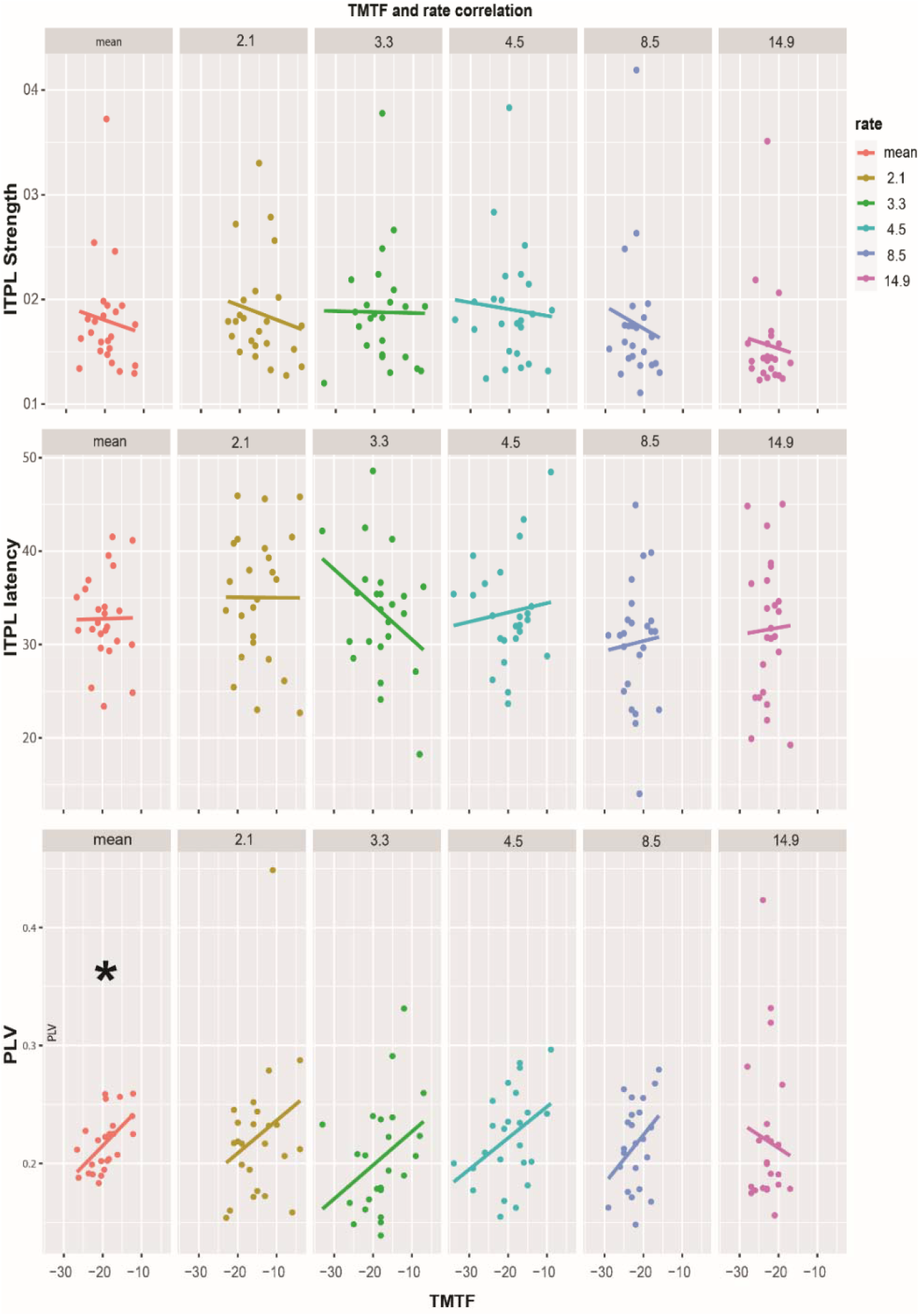
Brain-behavioral correlations for rate sensitivity. Behavioral TMTF thresholds did not show any association with neural *ITPL strength* nor *latency* at individual rates but for *PLV mean*. Scatters show brain-behavior correlations between TMTF and five different rates. We found no correlation with neural responses in ITPL. However, PLV pooled across rates (mean) showed a significant (^*^*p*<0.05) correlation.

## 4. Discussion

We compared cortico-acoustic tracking for speech vs. click stimuli across various speeds (below, within, and above the nominal speech rate) and levels of acoustic periodicity (jitter in 20% steps) to probe the effects of these stimulus factors on auditory cortex entrainment. By changing stimulus features and performing two types of phase-locking analysis (i.e., ITPL vs. PLV), we were able to parse the profiles of oscillatory activity coding sounds across trials (within tokens) vs. across time (between tokens).

For *rate*, we found token-wise oscillations showed a rightward hemispheric asymmetry in ITPL strength for both stimulus domains; stronger response at the nominal speech syllable rate (4.5 Hz) compared to faster rates; and longer response latency for speech vs. clicks. In contrast, phase-locking across tokens (i.e., PLV) to speech showed a surprising improvement in neural entrainment at 4.5 Hz but deteriorated at higher rates. For clicks, however, phase-locking strength dropped at 4.5 Hz and then rebounded sharply at 8.5 Hz until peaking at 14.9 Hz.

For *periodicity*, we found ITPL showed stronger phase locking in RH for both stimulus types but shorter latencies for clicks. Across token PLV, however, revealed that while phase-locking to speech declined with increasing jitter, entrainment to speech was still superior to that of clicks. In contrast to speech sounds, click responses were largely resistant to disruptions in periodicity. Collectively, our findings show that speech is overall more sensitive to changes in rate and periodicity and is also perhaps prioritized given the enhanced neural entrainment it evokes in the brain (even under passive listening).

### 4.1. Rate effects

Continuously changing speech rate/context requires ongoing adjustment by the listener (Casas et al., 2021). Our stimulus design included five rates intending to examine the auditory system’s temporal processing of speech and nonspeech sounds at speeds below, within, and above typical syllabic rates for speech (Poeppel et al., 2020). For both stimulus types, we found a phase-locking decrement with increasing rates, suggesting neural entrainment to periodic signals eventually collapses for very rapid sounds. Previous studies show neural oscillations align to rates higher than syllabic rates even at the expense of reduced intelligibility (Ahissar et al., 2001; Bosker et al., 2018). In this study, we found a rate effect in ITPL strength that was independent of stimulus type, suggesting neural entrainment (at the token level) is equally challenged for faster sounds regardless of their content, *per se* (i.e., speech vs. nonspeech). Gamma-band EEG responses have been linked to phonemic processing (Lizarazu et al., 2019) and higher-level post-perceptual processing (Di Liberto et al., 2015; Palva et al., 2002). However, similar rate declines for non-speech stimuli, and the passive design of our experiment indicate the need for broader explanations such as rhythmic (Pefkou et al., 2017), spectrotemporal (Giraud et al., 2012; Gross et al., 2013; Hyafil et al., 2015), or other such lower-level time-frequency coding and integration mechanisms.

#### 4.1.1 Domain specificity across rate (Speech vs. Click)

As with other auditory evoked responses, more transient signals (e.g., clicks) produce shorter response latencies than less transient signals (e.g., speech) due to their rapid onsets (Bidelman et al., 2018; Galbraith et al., 1990; Skoe et al., 2010; Song et al., 2006a). The longer onset ramping of our speech stimuli might explain the more prolonged response latencies we find for speech tokens compared to clicks. Additionally, speech requires more effort to process than clicks since it carries more complicated spectrotemporal patterns. The coding of speech and click tokens is thus differentiated in our token-level analysis not by their phase-locking strength, but by their timing (Song et al., 2006a), most likely as a result of differences in their signal bandwidth (Song et al., 2006b). The more gradual onsets of speech than clicks results in less synchronization and hence a longer latency (Kumar et al., 2011). Another explanation could be a greater degree of neural adaptation to longer speech than to faster click stimuli (Hoormann et al., 1992). With regard to the stimulus domain (speech vs. music), there was no distinction between hemispheres when the data were analyzed at the token level. This contrasts with the strong hemispheric effects we find in PLV when expanding the time window to longer stimulus segments and considering across-token neural entrainment (see *Section 4.1.3)*.

#### 4.1.2. Hemispheric laterality across rate

We showed a stark right hemisphere asymmetry in ITPL strength, regardless of stimulus type which is consistent with other studies (Casas et al., 2021). This rightward bias also confirms the “asymmetric sampling in time” hypothesis that emphasizes how the left auditory cortex is responsible for extracting information with a very rapid (∼ 20–40 ms) temporal integration window while the right hemisphere is biased for information at a slower (∼ 150–250 ms) temporal integration window (Ghitza, 2011; Poeppel, 2003; Zatorre et al., 2002). In other words, the right hemisphere is specialized for ∼4-6 Hz and the left hemisphere for ∼25-50 Hz. While we expected leftward laterality for speech, right lateralization has also been observed in previous experiments utilizing speech stimuli (Abrams et al., 2008; Alexandrou et al., 2017; Gross et al., 2013). However, given the fact that all rates used in this experiment were less than ∼25 Hz, an interesting finding in our data is the same RH asymmetry with even click (non-speech) stimuli. This emphasizes that the cerebral hemispheres are perhaps specialized based on the speed (rate) of auditory information rather than the stimulus domain, at least under the passive listening conditions of our task. Latency also demonstrated a rate x hemisphere interaction, indicating that the speed of neural response elicitation depends upon hemisphere and rate. Therefore, our prediction to see different hemisphere activity for each stimulus type was not validated since all neural oscillatory activities were tuned to rates below 25 Hz.

#### 4.1.3. Within vs. between token phase-locking across rate (ITPL vs. PLV)

In contrast to ITPL, we found an interaction between rate and stimulus type for PLV measures. ITPL denotes phase-locking to a stimulus on a trial-by-trial basis. As a measure of evoked response consistency, it examines the reproducibility of neural activity across individual trials of a token. As such, the temporal integration window of our ITPL analysis was relatively short as it only considered neuronal responses occurring within 60 ms of stimulus onset. In terms of speech, this limited window represents the phonemic level of language hierarchy. Gamma oscillations (like those in our 30-40 Hz analysis band of the ITPL spectrograms) are typically nested or phase-coupled within lower frequency theta waves (Giraud et al., 2012; Peelle et al., 2012). Slower theta waves easily follow the envelope of ongoing speech and are thus thought to integrate syllable representations across tokens—a larger level of the speech-analysis hierarchy (Casas et al., 2021). Thus, the lack of distinguishment between speech and click entrainment in ITPL might have resulted from the much small processing window and consequently limited timescales of oscillatory activity that can develop at the token level. In other words, 60 ms of temporal integration seems insufficient to differentiate clicks from a speech in terms of auditory entrainment; both require many seconds before the brain can integrate and differentiate signal content, speech or otherwise (Roß et al., 2002). Of course, this is only in terms of entrainment. Speech and clicks are otherwise easily differentiated in only a few milliseconds of the signal for dimple identification tasks (Knebel et al., 2018).

In contrast to within-token ITPL, we found PLV strongly differentiated speech vs. non-speech stimuli. PLV reflects the phase-locking between the acoustic stimulus and EEG response over the entire course of the stimulus stream. Therefore, PLV is a measure of the robustness of brain synchronization *across tokens*, and thus captures oscillatory activity at a higher level of analysis [cf. sentence (PLV) vs. phonemic (ITPL) level of processing]. Surprisingly, across-token PLV to speech revealed strong enhancements in phase-locking at 4.5 Hz, validating the concept of a universal speech syllabic rate across global languages (Assaneo et al., 2018). Critically, this enhancement at ∼4.5 Hz was specific to speech and did not occur for clicks (i.e., domain-specific effect). Moreover, we found phase-locking to speech plummeted at higher frequencies, suggesting these speech-specific enhancements are limited to stimulus rates that occur in natural for speech (Assaneo et al., 2018). At higher rates, and in contrast to speech, click synchronization actually improved. The counterintuitive enhancement of click responses at the higher rates is however consistent with psychoacoustical findings (including those here), explained as increased summation (i.e., temporal integration) of postsynaptic potentials as a result of activating a broader neural population (Brugge et al., 2009; Forss et al., 1993). Thus, while the auditory system is certainly capable of synchronizing to higher stimulus rates (as evidenced by our click data), it appears as though the sensitivity to modulations in speech synchronization is more restricted.

### 4.2. Periodicity effects

Various temporal scales of the speech signal and linguistic hierarchy are correlated with neural entrainment (Ghitza et al., 2009). Given that speech is not perfectly periodic and shorter syllables frequently follow or precede longer ones, it is intriguing to examine how the periodicity/aperiodicity of speech influences brain entrainment (Ghitza et al., 2009). By parametrically varying the jitter of otherwise periodic signals, we aimed to perturb the input integrity and examine how brain rhythms to nominal speech rate (at 4.5 Hz) are disrupted by aperiodicity. Our results reveal minimal jitter effects in ITPL strength and latency, suggesting token-level coding is largely impervious to aperiodicity. In classic evoked response paradigms, which only consider token-wise responses, the temporal information (e.g., interstimulus interval) between successive stimulus events does not directly affect the neural representation of the acoustic signal, at least at lower rates where adaptation would play an effect. However, other studies showed that “cross-token comparisons” could be a cue for temporal coding that has a significant impact on intelligibility (Huggins, 1975; Miller et al., 1950). Thus, temporal perturbation can affect how sounds are organized and subsequently perceived. Our PLV measurements robustly detect these jitter effects, which occur over a longer time frame, especially for speech events.

Periodicity in speech can facilitate perception and intelligibility (Benesty et al., 2008). Ghitza et al. (2009) altered the interstimulus intervals of syllables using periodic and aperiodic interruption in order to affect intelligibility and ascribed neural entrainment to internal processing rather than acoustics aspects of the sounds (Ghitza et al., 2009). They showed the speech intelligibility of the compressed signal is poorer to that of the original signal. In their study, inserting 20-120 millisecond-long gaps of silence into a speech significantly increases its intelligibility. Our study design excluded an intelligibility component as it was passive and focused arguably on only the acoustic-phonetic characteristics of speech. Still, our results reveal that neural entrainment is improved for rhythmic speech as opposed to click or even aperiodic speech. It is possible such neural effects account for the perceptual facilitation observed for periodic signals in previous behavioral studies (e.g., Ghitza et al., 2009).

#### 4.2.1. Hemispheric laterality

Only the 4.5 Hz presentation rate was chosen in the jitter experiment, as it was hypothesized to play the most significant role in the rate effect (Assaneo et al., 2018). According to one theory, lateralization of neuronal processing in hearing is induced by the temporal pattern of auditory perception and cortical asymmetries by underlying disparities in neural populations with differential temporal sensitivity (Poeppel, 2003; Zatorre et al., 2002). This cortical cytoarchitectonic difference suggests that the right hemisphere more precisely tracks modulation frequencies within the syllable rate range (Assaneo et al., 2019a) confirming our findings in ITPL strength. To the best of our knowledge, this is the first study that describes lateralization in various periodicities and we could not find other evidence for hemispheric differences in periodicity sensitivity. Since the same pattern of hemispheric effect was observed in both rate and periodicity, we attribute our findings to the effects of temporal integration windows, as describe earlier (Ghitza, 2011; Poeppel, 2003; Zatorre et al., 2002).

#### 4.2.2. Within vs. between tokens

Our PLV data demonstrate periodicity did not affect across-token phase locking to clicks. Despite the fact that clicks are more perceptually salient than speech due to their rapid onset (Vigil et al., 2020), jitter did not affect PLV entrainment sensitivity for clicks. With increasing the jitter of speech stimuli, however, phase-locking deteriorated. It is possible that as speech becomes more aperiodic, the brain treats the signal more like a non-speech stimulus, resulting in similarly low PLV as we observe for click stimuli. This could explain why stimulus type and jitter interacted in our PLV analysis. Periodicity in speech may boost neural responses via predictive coding, which could account for the stronger entrainment to speech we find at low relative to high degrees of jitter (Peelle et al., 2012; Thut et al., 2011). Previous research also reveals specific populations of neurons that respond to aperiodic but not periodic stimuli (Yrttiaho et al., 2008). Also, imaging studies demonstrate the location of periodicity-sensitive cortical areas can be more anterior than aperiodic stimuli, depending on the stimulus type (Hall et al., 2009). Our technique for EEG source analysis only localized responses in the auditory cortex, which may not elicit equivalent responses for speech and click stimuli.

### 4.3. Brain-behavior relations between entrainment and rate/periodicity sensitivity

We demonstrate an association between behavioral sensitivity and brain synchronization to stimuli varying in rates. Individuals who showed greater sensitivity in psychoacoustic tests (behavioral responses) exhibit less robustness between token measures. A recent study by Casas et al. (2021) also showed that participants that are more behaviorally responsive to temporal changes in ongoing sounds exhibit weaker phase-locked responses. They attributed their counterintuitive findings to the sample size and other methodological concerns including in the perceptual difficulty of their stimulus set (Casas et al., 2021). However, one possibility is that their neural recordings included several entrained responses from multiple brain areas (not exclusively auditory regions). Indeed, conventional EEG suffers from volume conduction resulting in frontal generators contributing to auditory evoked responses (Bidelman, 2016; Knight et al., 1989; Picton et al., 1999). Our use of passively presented stimuli and source analysis helps exclude attentional or task confounds (as suggested by Casas et al) that are likely modulated by frontal cortical areas. Having the same result, however, it is possible that individuals who show better brain-to-acoustic coupling relegate temporal processing to lower levels of the auditory system (e.g., brainstem, thalamus), more peripheral to the behavior and cortical responses assessed here. Whereas others who perform worse or more laboriously in temporal processing tasks might require high levels of auditory processing at the cortical level as a form of compensatory mechanism (Momtaz et al., 2021; Momtaz et al., 2022). This might account for the counterintuitive negative correlation we find between cortical phase-locking strength and behavior, i.e., more effortful encoding (higher PLV) in less perceptually sensitive individuals.

One significant distinction between our brain and behavior evaluations is that although TMTFs measure rate modulation thresholds in active detection, neural recordings were under strictly passive listening. It is highly likely the nature of the correlation would change had we conducted our behavioral tasks during the EEG recordings. Similar arguments could be made for the CA-BAT, which measured periodicity threshold.

## 5. Conclusions

Overall, this study aimed to address questions regarding rate, periodicity, and stimulus differences in EEG neural entrainment and their association with behavioral responses. By examining the same parameters from two distinct perspectives (ITPL vs. PLV analysis), our data reveal unique distinctions in how each of these factors impacts the neural encoding of ongoing complex sounds. A summary of the major findings of our study is provided **Table 1**.

**Table 1.**
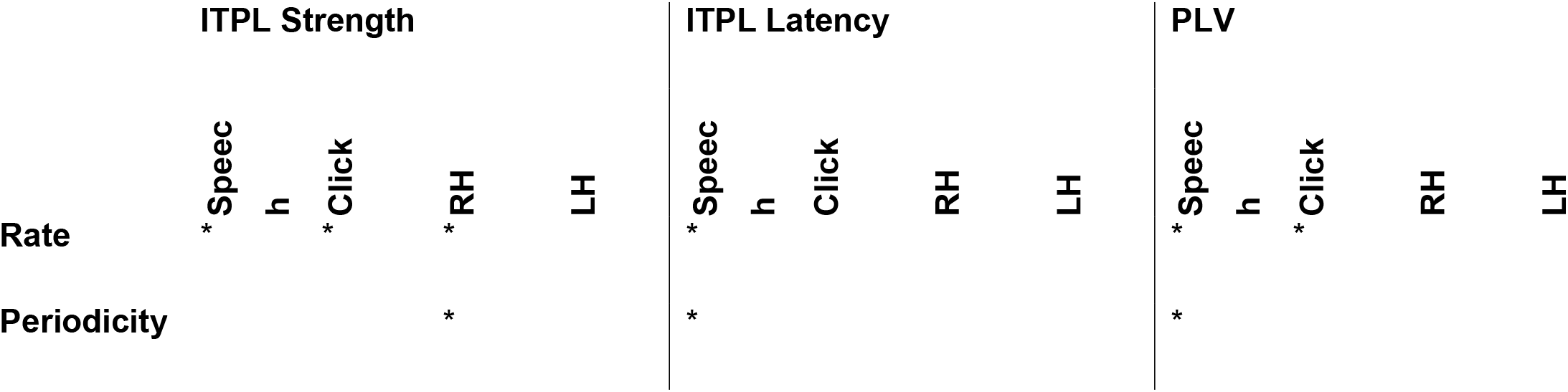
Summary of significant rate and jitter effects for the different dependent variables across the stimulus manipulations that were studied. * = significant parameters.

Having revealed speech is particularly rate and periodicity sensitive (more so than non-speech clicks), this might inform broader work examining temporal processing issues in patient populations such as those with certain auditory processing disorders (Momtaz et al., 2021; Momtaz et al., 2022) or dyslexia (Ben-Yehudah et al., 2004; Tallal et al., 1993) which impact auditory temporal processing. The data here characterize the constraints of temporal capabilities in auditory cortex and neural entrainment. It would be interesting to extend the current paradigm to future studies in these clinical populations. Auditory plasticity induced by training and rehabilitative programs that aim to enhance temporal processing (Anderson et al., 2013; Chermak et al., 2002; Moreno et al., 2014) could be used to enhance cortical phase-locking and, subsequently, speech understanding in challenging listening environments. The stimulus specificity we observe in entrainment patterns also speaks to the need to incorporate the correct stimuli in training plans if the goal is to maximize neural and behavioral outcomes. Indeed, our data suggest periodic speech at or near nominal syllabic rates (4.5 Hz) might have the largest impact on perception and cognition following rehabilitation. Future studies are needed to test these possibilities.

The data here demonstrate that periodic speech differentially affects neuronal entrainment in a passive condition; however, how mechanisms might change in an active attentional setting remains unknown. Presumably, active tasks during entraining stimuli might also recruit additional (non-auditory) brain regions other than the auditory cortex, which was the focus of this investigation. In this vein, functional connectivity approaches (Rimmele et al., 2018) might be used to further tease out the dynamics of bottom-up (sensory-driven) versus top-down (cognition-driven) processing and interhemispheric connections that affect the brain’s ability to entrain to rapid auditory stimuli.

## Acknowledgments

This work was supported by grant NIH/NIDCD R01DC016267 awarded to G.M.B. The authors thank Dr. Eugene Buder for his comments on earlier version of this manuscript.

## Footnotes

1 Note this definition is similar to ITPL (Eq. 1) with the exception that phase-locking across tokens is computed over time (sample-to-sample) rather than across trials of the stimuli presentation as in Eq. 1. The terms ITPL and PLV are otherwise interchanged and both reflect the degree of phase consistency between neural and stimulus signals. Here, we use “ITPL” and “PLV” to refer to oscillations analyzed within-vs. between-tokens, respectively.

